# An interplay between cytoplasmic flow and cell morphology during cytokinesis simulated by the moving-particle semi-implicit method

**DOI:** 10.64898/2025.12.22.695877

**Authors:** Yusuke Yoshimi, Yuki Tsukada, Noriko Hiroi, Akatsuki Kimura, Akira Funahashi

## Abstract

Cells are composite mechanical systems: a viscoelastic cytoplasm enclosed by a deformable cell cortex. These two components exchange forces continuously—fluid motion deforms the cortex, and cortical deformation in turn redirects intracellular flow. Yet most cellular mechanics simulations treat either the fluid or the cortex in isolation, leaving their mechanical interplay largely unexplored. To address this gap, we developed a fluid–structure interaction framework based on the moving-particle semi-implicit (MPS) method that solves the coupled dynamics of cytoplasmic flow, internal pressure, and cortical elasticity. We applied this framework to cytokinesis, focusing on how the cleavage plane is positioned. While the mitotic spindle is known to specify the furrow site, prior work has emphasized biochemical cues emanating from the spindle, whereas a direct mechanical cue has been difficult to interrogate. Our simulations show that spindle elongation generates cytoplasmic flows that create a pressure minimum near the spindle midzone; when the cortex is modeled as an elastic shell, this pressure landscape drives a localized cortical invagination at that site. We therefore propose that spindle-driven cytoplasmic flow provides a mechanical positional cue for cleavage-furrow initiation.

**Author Summary:** Dividing cells exhibit two contrasting properties: dynamic state changes and the maintenance of homeostasis. During cell division, a dividing cell breaks its identity while successful daughter cells inherit a certain set of its components for survival. Heterogeneous physical properties such as cell shape, cytoplasmic flow, and internal pressure interact to maintain an appropriate range of cellular conditions. The interaction of such physical components is highly dynamic; therefore, simulation studies play an important role in understanding the mechanisms. We constructed a model based on the moving-particle semi-implicit method that recapitulates the behavior of highly dynamic physical objects such as cytoplasmic fluid with practical computational time. The simulation showed that the change in cell shape is caused by internal cell pressure, which is evoked by cytoplasmic flow during elongation of spindle microtubules. The developed simulation platform provided a tool to integrate physical measurements to understand the mechanisms that balance physical dynamics during cell division. Such quantitative simulation combined with the data from molecular or genetic experiments may predict how internal dynamics maintain homeostasis during drastic changes in the cell state.

## Introduction

Cytokinesis is a highly dynamic process that visually begins with formation of a cleavage furrow on the previously uniform cell surface. Before the cleavage furrow formation, the spindle apparatus is rearranged in preparation for cell division [1]. The spindle elongates, and the contractile ring is formed with actin and myosin at the division plane. Regulatory signaling molecules such as Rho GTPases orchestrate these cytoskeletal dynamics [2]. Although the details of each molecular process involved in cytokinesis have been uncovered by genetic and molecular studies, the interactions among different mechanisms remain unclear [3]. It is currently debated which mechanism determines the plane of cytoplasmic division. Spindle reorganization and elongation occur during anaphase, and elongation continues during cytokinesis to form the division plane; the spindle and its microtubules play an important role in determining the position of the division plane [4]. Some molecular mechanisms have been proposed by which astral microtubules and the central spindle regulate the localization of signaling molecules that determine the division plane [5]. In this study, we focused on cytoplasmic flow generated by spindle elongation as a potential mechanism to shape the cell division plane at the appropriate position.

Cytoplasmic flow regulates the dynamics of signaling molecules and of the cytoskeleton, such as actin filaments. White and Borisy have proposed a mechanism in which cytoplasmic cortical flow accumulates the actin–myosin network on the cell equatorial belt and causes tension to induce cytoplasmic division [6]. Reymann et al. have quantified the relationship between cytoplasmic cortical flow and the alignment of actin filaments around the equator during the formation of the contractile ring [7]. Thus, the cytoplasmic cortical flow may align polarized actin filaments to induce the contractile ring during cytokinesis [7–9]. Cytoplasmic cortical flow transports molecules and contributes to the localization of cytoskeleton regulators to induce formation of a cell division plane [10]. We hypothesized that the rheological influence of cytoplasmic flow is also a considerable factor in understanding the regulation of cell division plane formation. Inward cell cortex depression may be caused by a localized decrease in internal pressure. Conversely, a localized increase in pressure may induce cell surface extension by interacting with the elasticity of the cell cortex. Investigation of these physical interactions is essential to understanding the dynamics of cell division.

To examine the regulation of cell division by cytoplasmic flow from the rheological point of view, a mechanism of flow generation should be considered. We postulated a mechanism of flow generation by spindle extension during anaphase. In *Caenorhabditis elegans*, cell division can be inhibited by the laser ablation of the spindle microtubules [11] or by the genetic disruption of the spindle microtubules [12]. The cytoplasm enters the increasing space generated by spindle elongation, consequently generating cytoplasmic flow through the entire cell. Thus, spindle elongation may drive cytokinesis through cytoplasmic flow generation.

Fluid dynamic analysis is suitable for examining the relation of internal pressure and cytoplasmic flow [13]. To test the hypothesis that the cell division plane is determined by cytoplasmic flow, we performed fluid dynamics analysis together with computational simulations that model cytoplasmic flow generated by spindle elongation. Computational fluid dynamics is a research field that provides tools for understanding the principles of spatio-temporal dynamics of the fluids. Applications of computational fluid dynamics cover studies of mitotic spindle positioning [14] and cell division [15], and serve as a basis for describing the interaction between the whole body and surrounding environment [16]. Technical challenges of rigorous computational fluid simulation applied to cell division include maintaining the balance between accuracy and computational cost and taking into account the dynamics of the different physical components in the target cell. Cytoplasmic flow and internal pressure should be integrated to model the behavior of the entire cell. Moving particle semi-implicit (MPS) simulation based on the implicit method is known to be efficient in terms of stability and computational cost [17].

Here, we addressed the mechanism that determines the position of the cell division plane during cytokinesis based on the fluid dynamic model for the cytoplasmic flow caused by spindle elongation. We constructed and analyzed a computational simulation model to investigate the interaction between cytoplasmic flow and the elastic cell cortex. The model includes the generation of cytoplasmic flow by spindle elongation, a description of the internal pressure generated by the cytoplasmic flow, and cell cortex morphology. Simulation showed the regulation of the cell morphology dynamics in a self-organizing manner through cytoplasmic flow. We assessed the consistency between numerical experiments and experimental quantification of the fluorescence time-lapse images of the labeled centrosome and cell morphology in *C. elegans* embryonic cells. We also conducted simulations using an arbitrary triangular cell shape to validate against experimental results of triangular cell shape.

## Results

### Cell morphology simulation with the MPS method

We conducted MPS simulations of cytoplasmic flow and cell shape deformation during the first division of early *C. elegans* embryos. The model consists of spindle microtubes, centrosomes, the cell cortex as an elastic cell surface, and the fluid that comprise cytoplasmic flow (Fig. 1). The simulation describes physical interactions among the spindle dynamics, cytoplasmic flow, and the cell cortex elasticity. Spindle elongation induces cytoplasmic flow by increasing the local volume, and the flow modulates pressure toward the cell cortex. Consequently, our simulation draws the dynamics of cell shape change. Simulation parameters are shown in Table 1. The simulation begins with the elongation of the spindle during anaphase, since we focus on the determination of the cell division plane during telophase. The speed of the spindle elongation v = 5.0 × 10^−8^ m/s was adopted from [18]. In the model, the spindle is defined as a sphere formed from two centrosomes and microtubules extending from it; its size estimation (7.0 μm) was based on the fluorescence image of the spindle. We followed the model proposed in [19] that considered the cytoplasmic flow change as a consequence of microtubule movement. Our computational model simulates the dynamics of cytoplasmic flow, cell internal pressure, force at the cell cortex as the outcome of the spindle elongation, and interaction among them.

**Fig 1.**
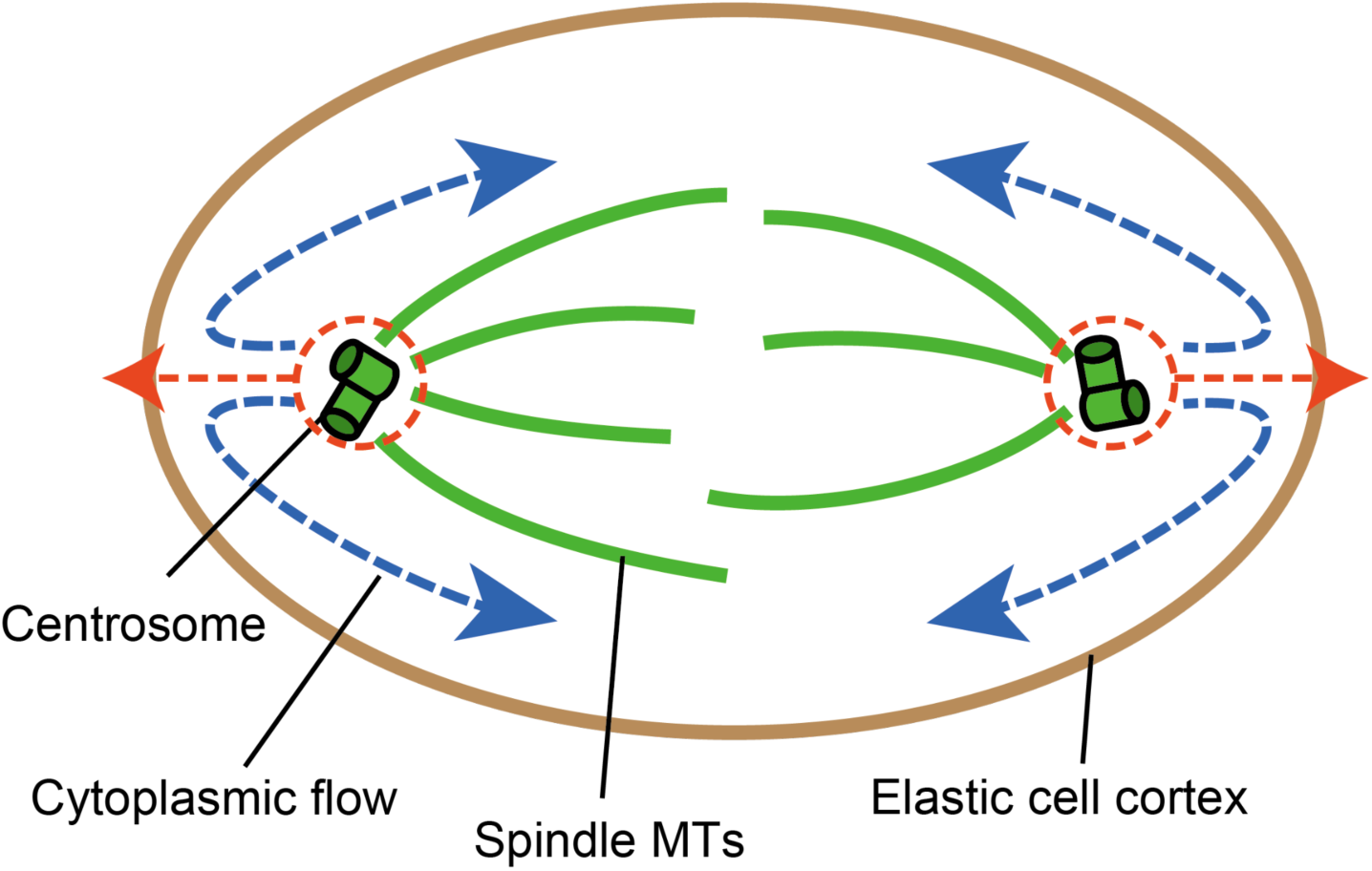
Scheme of the mathematical model of cytokinesis. The components of the model are labeled; MTs, microtubules. Each component has different physical properties and interacts with the other components. Red arrows, centrosome movement as the consequence of spindle MT elongation. Blue arrows, cytoplasmic flow generated by the centrosome movement. Under internal pressure related by the cytoplasmic flow, the cytoplasmic flow interacts with the elastic cell cortex.

**Table 1.**

| Parameter | Value | Reference or remark |
| --- | --- | --- |
| Viscosity $\mu$ | 1.0 (Ns/m) | *C. elegans* embryo [20] |
| Density $\rho$ | 1000 (kg/m<sup>3</sup>) | Water density |
| Kinematic viscosity coefficient<br>$\nu = \mu/\rho$ | 1 (mm<sup>2</sup>/s) | - |
| Length of the cell along the major axis | 50 (μm) | - |
| Length of the cell along the minor axis | 30 ($\mu\text{m}$) | - |
| Spindle radius | 50 (nm) | *C. elegans* embryo [18] |
| Elongation speed of the spindle | 50 (nm/s) | *C. elegans* embryo [18] |
| Spring constant in the direction of cell<br>cortex elongation | 1 ($\mu\text{N/m}$) | Red blood cell [21] |
| Spring constant in the direction of cell<br>cortex bending | 1 (pN/m) | Red blood cell [21] |

### The dynamics of cytoplasmic flow

We performed a fluid simulation from the start of spindle elongation (t = 0 s) to its stop (t = 200 s) and visualized cytoplasmic flow. The input elongation time series for the simulation is shown in Figure 2A. Using MPS based on Navier–Stokes equations, our model computed the constantly elongating spindle that pushed the cell cortex represented by the elastic surface and the generation of cytoplasmic flow. The simulation generated the internal direction of cytoplasmic flow on the cell cortex, which followed spindle elongation during cytokinesis (Fig. 2B, C); the flow was generated promptly after starting the simulation, and distinct patterns were maintained until spindle elongation slowed down (t = 150 s). At around t = 200 s, the elongation speed was almost zero, and the distribution of the flow directions became random. During the emergence of distinct cortical flow appeared at t = 2 s, the boundary between the blue and red regions in Fig. 2B, C denotes where the opposite streams meet. The boundary corresponds to the middle of the spindle, which is illustrated by the dotted black line where the contractile ring appears. This simulation supported the hypothesis that cortical flow accumulated materials that comprise the contractile ring.

**Fig 2.**
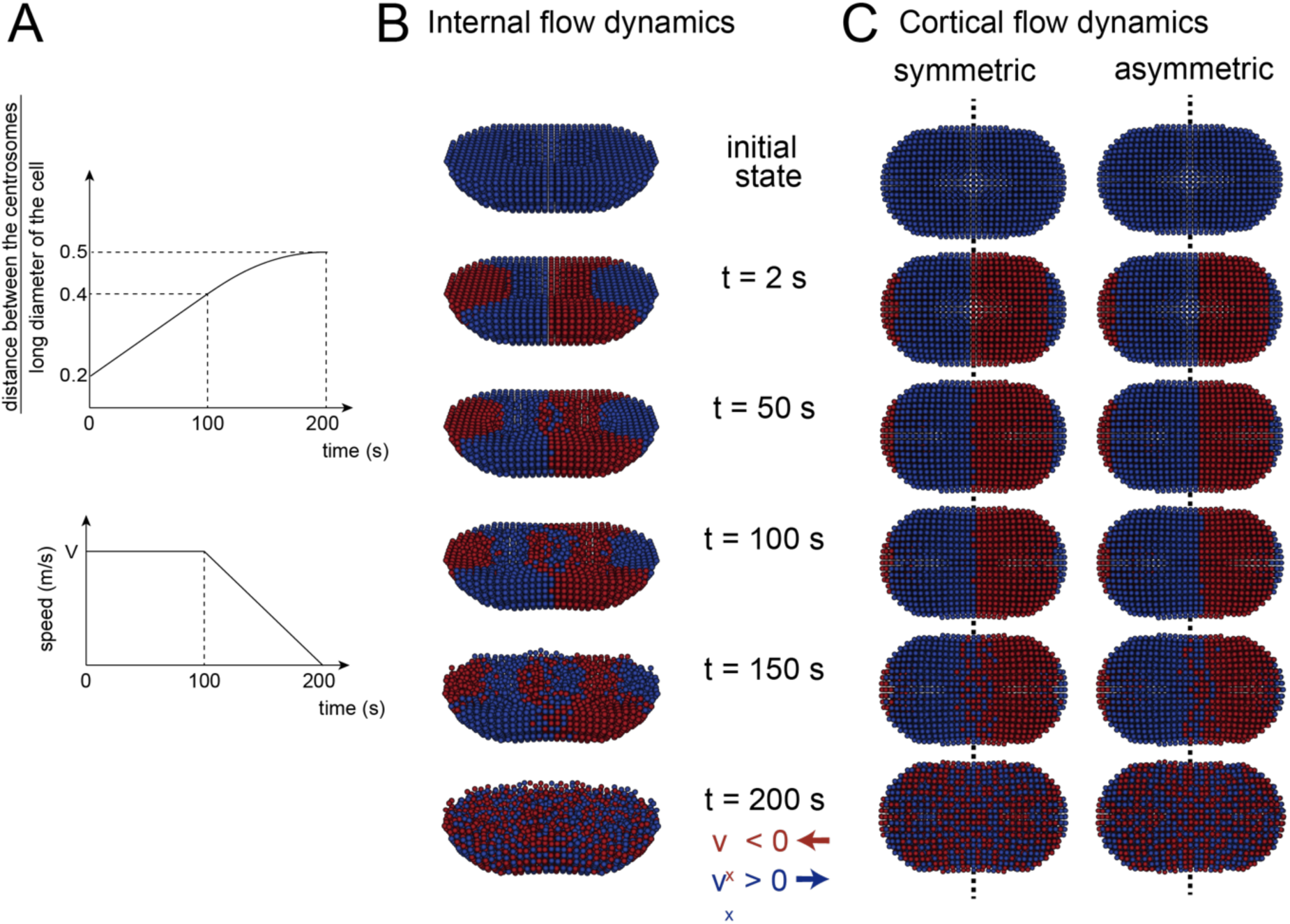
Simulated cytoplasmic flow. (**A)** The input spindle elongation time series for the simulation. (**B)** Simulated cytoplasmic dynamics. Only the bottom hemisphere is shown to depict the cell’s internal conditions. Colors indicate positive (blue) and negative (red) velocity along the cell’s major axis. **(C)** Simulated cytoplasmic dynamics on the cell cortex. Surface velocity is shown similar to (B). The dotted black line illustrated the middle of the spindle.

In asymmetric cell division, spindle asymmetry precedes the asymmetry of cell shape, and a contractile ring forms in the middle of the asymmetric spindle [22]. To test whether the boundary position for the opposite directional flow follows the asymmetric localization of the spindle, we performed a simulation by shifting the initial position of the spindle to the posterior side (Figs. 2C); the spindle elongation speed was the same. The boundary of the opposite directional cytoplasmic flow was formed in the middle of the spindle, not at the center of the cell.

### Internal pressure distribution in the cell caused by cytoplasmic flow

Cytoplasmic flow affects the rheological balance inside the cell, resulting in a nonuniform internal pressure distribution. A simulated time course of pressure distribution generated by cytoplasmic flow during cytokinesis is shown in Figure 3. At around t = 2–100 s, pressure was increased at both of the spindle poles and decreased in the middle of the spindle. In asymmetric cell division, pressure decreased in the middle of the spindle, not at the center of the cell. Pressure distribution was nonuniform at t = 200 s.

**Fig 3.**
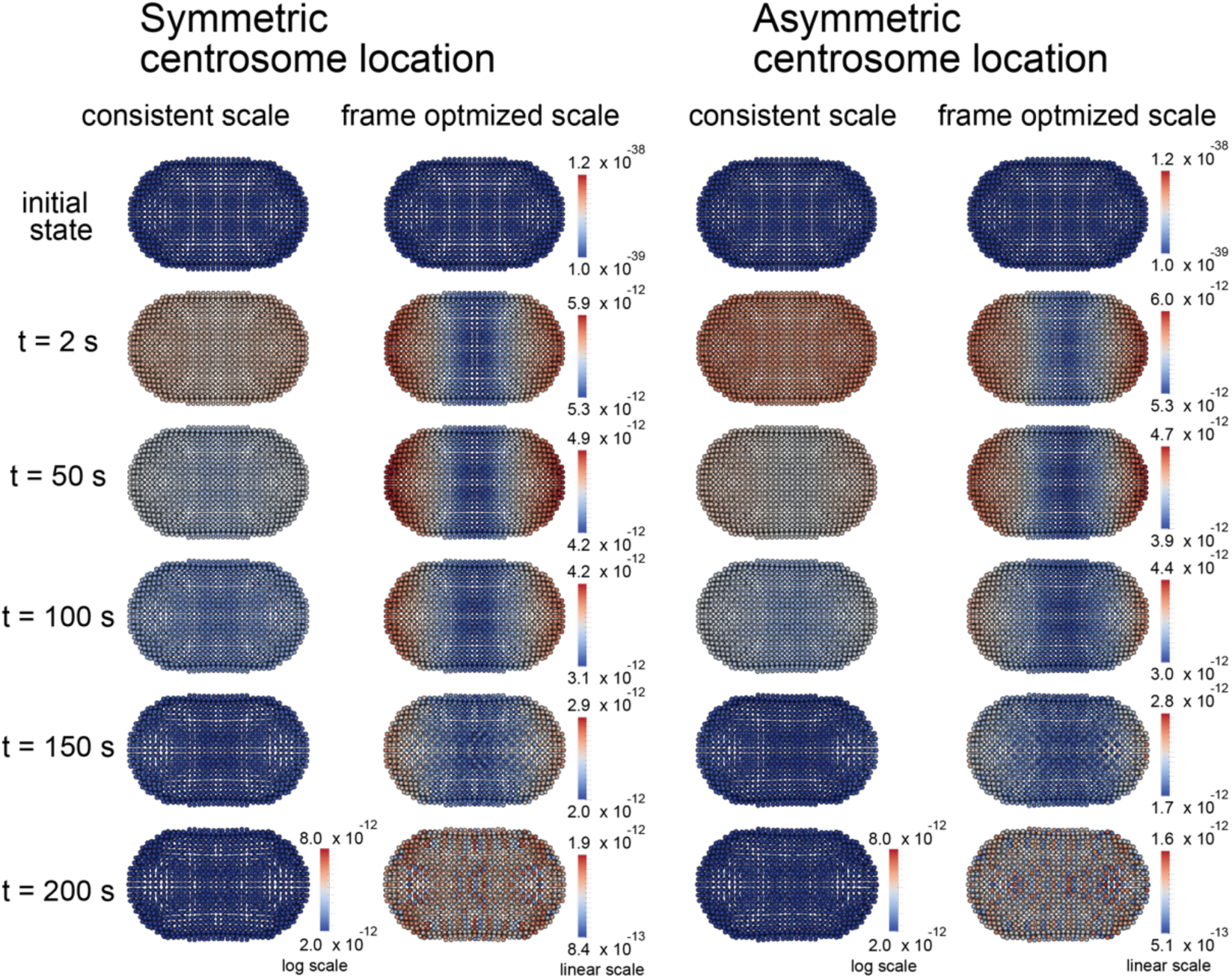
Simulated distribution of pressure on the cell cortex. Symmetrical and asymmetrical centrosome locations are set as the initial condition. Each result is displayed according to a log scale color bar adjusted throughout the entire time course, or on a linear scale color bar with adjustments in each time frame.

### Cell deformation as the outcome of cytoplasmic flow

The deformation of the cell surface induced by the interaction between the internal pressure distribution and the elastic cell cortex during cytokinesis is shown in Figure 4. Cytoplasmic flow induced internal pressure and consequently evoked a concavity at the cell surface on the plane of the center of the spindle poles, which became apparent around t = 100 s (Fig. 4A). Closely extracted lateral cortex shape showed an emergence of concavity around t = 50 s (Fig. 4C) following pressure change shown in Figure 3. The position of the concavity is determined by the center of the spindle poles. The simulated change in cell shape is derived from the computation of the rheological interaction between the elastic cell cortex and internal pressure caused by cytoplasmic flow, and occurs in a self-organizing manner. At t = 200 s, pressure distribution became mosaic (Fig. 3), but the concavity remained at the same position. Thus, cell morphology did not immediately follow the internal pressure distribution, but the cell’s internal kinetics maintained the deformation. We assume that the small morphological modulation triggers the localization of signaling molecules, such as the small GTPases involved in cytoskeletal regulation as proposed in previous reports [23,24], and induces the generation of the contractile ring as an initial determinant for the cell division plane [25].

**Fig 4.**
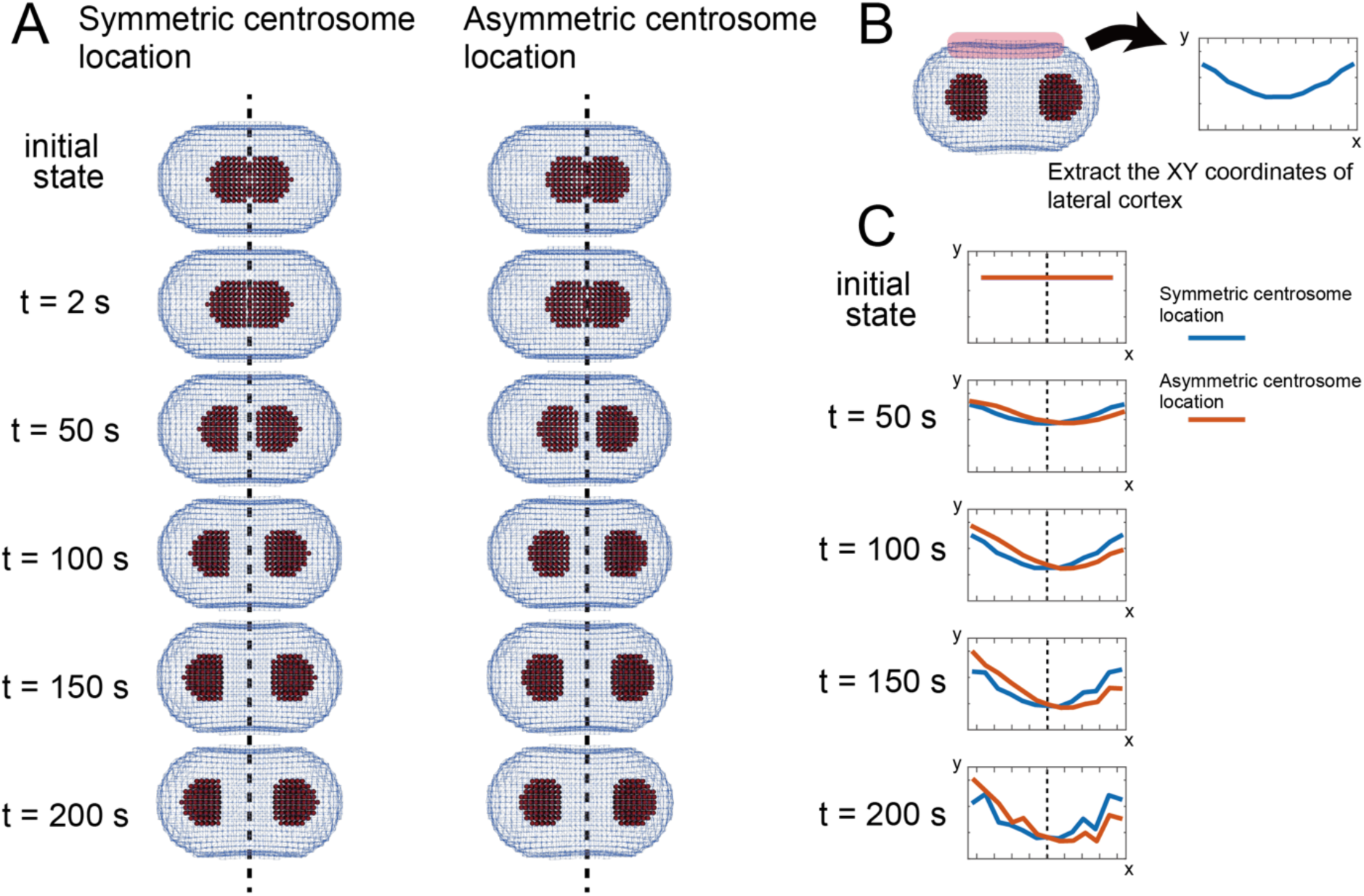
Time course of the cell morphology change. **(A)** Snapshots of the simulated cell shape drawing with a wireframe expression. Red particles, centrosomes. Dashed lines, the initial midline of the centrosomes. The emerging concavity is located around this initial midline. (**B)** The *xy* coordinates of the lateral cortex particles at the center of the cell along the z-axis are visualized to clarify morphological changes during the simulation. **(C)** The extracted positions of lateral cortex particles. Dashed lines show the initial midline of the centrosomes.

### The concavity position is independent of the decrease in spindle elongation speed

Our simulation in Figure 4 showed that the concavity position induced by cytoplasmic flow is located at the equatorial plane of the spindle regardless of where the spindle is located; a concavity position is determined by the center of the spindle. This result evokes a question of whether the elongation dynamics of the spindle microtubules affect the internal pressure distribution and the position of the induced concavity. To address this question, we conducted simulation with a perturbation of spindle elongation speed by setting slow elongation speed (3.75 × 10^−8^ m/s) until the end of the simulation. At both control and perturbated conditions, a concavity was located at the middle of the spindle (Fig. 5). To clarify the time course of the cell morphology change, we calculated the curvature of the cell surface around the contractile ring. The curvature *κ* was defined by the cell boundary curvature along the major axis of the cell at *z* = 0 (Fig. 5B). We then plotted the time course of the normalized maximum curvature of the extracted profile (Fig. 5C). Decreased speed delayed curvature increase, supporting the link between spindle elongation and furrow formation.

**Fig 5.**
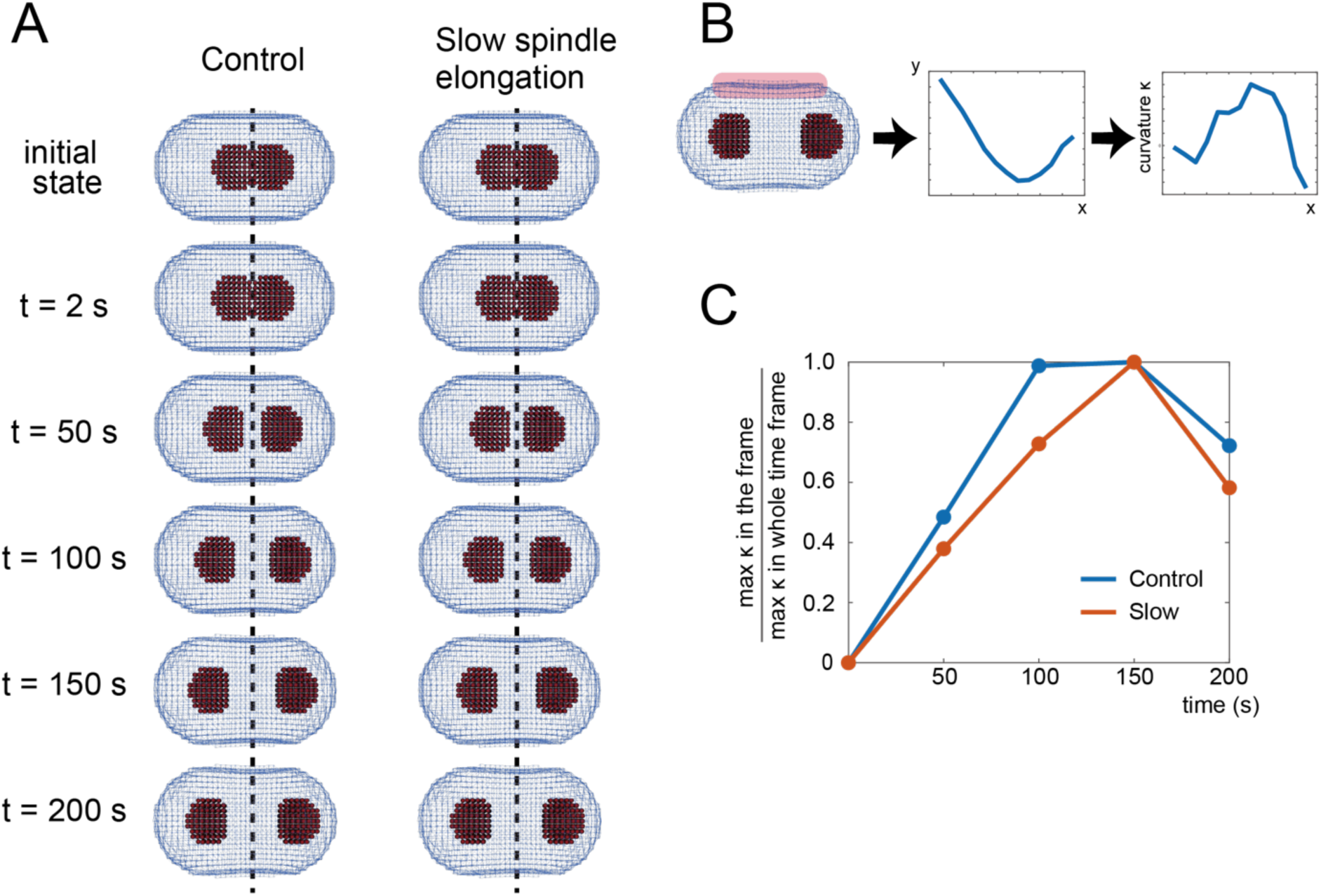
Simulation of the reduction of the spindle elongation. **(A)** Comparison between the simulations of control and slow spindle elongation conditions. Centrosomes are initially asymmetrically located. There is no clear difference between the control and slow spindle elongation in the position of the invagination at t = 200 s. The dashed lines indicate the initial midline of centrosomes. (**B)** The *xy* coordinates of the lateral cortex particles were extracted as in Fig. 4, and then the curvature along the *x*-axis was calculated. **(C)** Time courses of the normalized maximum curvature at each time point. Slow spindle elongation delayed the curvature change and a quick decrease of the curvature at t = 200 s.

### Quantitative image analysis for elongation-defective embryos (*gpr-1*/*gpr-2* RNAi)

We experimentally tested our model’s prediction on a delayed onset of the furrow with slow speed of spindle elongation. Elongation is positively regulated by GPR-1 with the G-protein alpha-subunit–binding domain and GPR-2 with the GDP-dissociation inhibitor domain [26] supported by double RNAi experiments of two related G *α* _i/o_ subunits, *goa-1 and gpa-16,* causes spindle defects that are indistinguishable from *gpr-1/gpr-1(RNAi)*[27]. We conducted RNAi for *gpr-1/gpr-2* in the CAL0092 strain, which carries GFP-labeled gamma-tubulin for centrosome visualization. Using the time-lapse images of fluorescence microscopy, we measured the time courses of the distance between centrosomes and cell curvature during cytokinesis.

We first compared the spindle elongation speed of the control and *gpr-1/gpr-2* knockdowns. The time courses of the distance between the spindle poles are shown in Figure 6A. In the control, a pronounced increase in the distance was observed at t = 200–400 s, whereas the timing of the separation was delayed and the speed was decreased in *gpr-1*/*gpr-2* knockdowns. The average elongation speed was lower in the knockdowns than in the wild type, and the final distance between the spindle poles became shorter.

**Fig 6.**
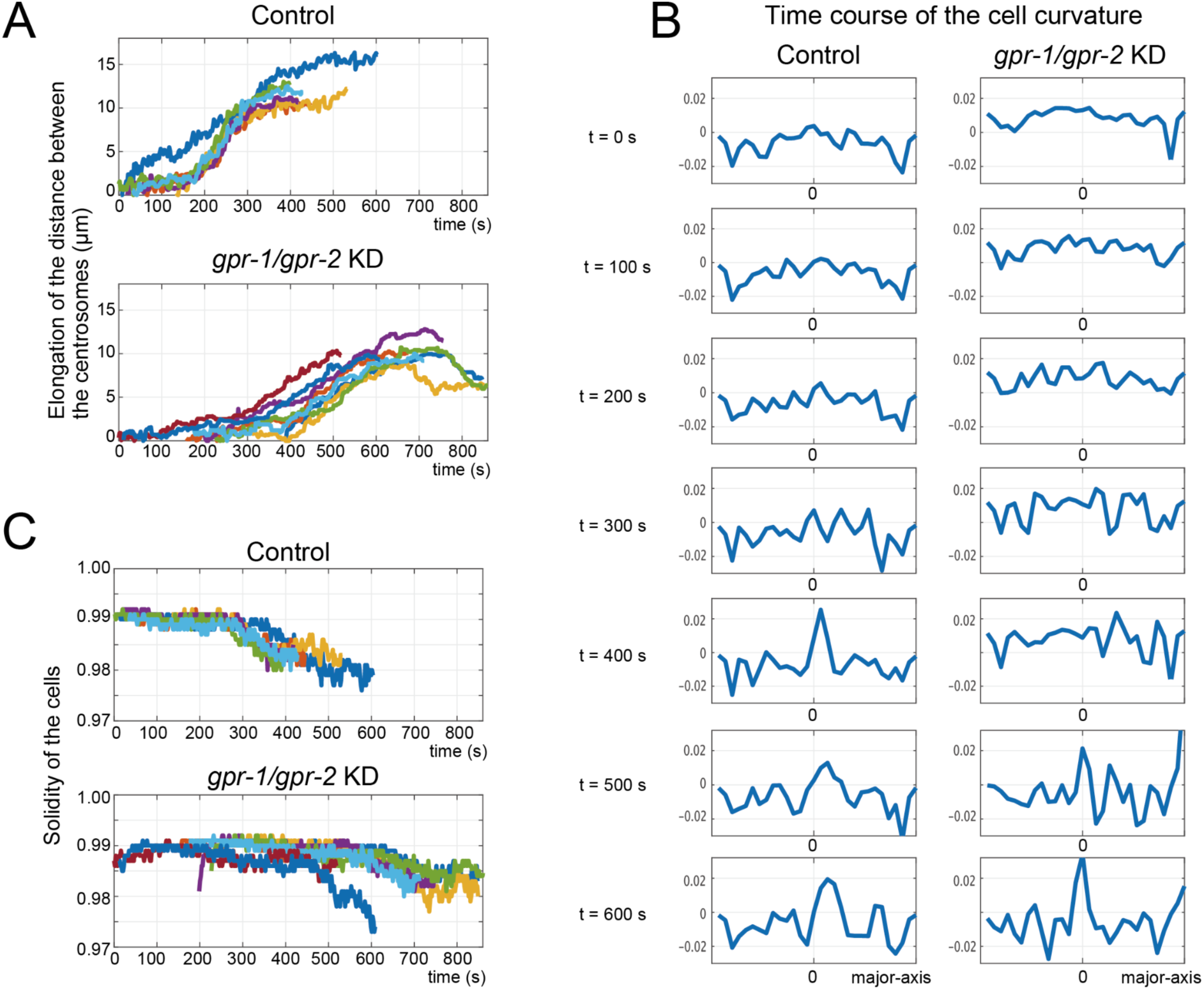
Quantification of time-lapse fluorescence images. **(A)** Time courses of the distance between centrosomes. t = 0 s is the time of NEBD. The initial distance was set to 0 by subtracting the initial distance from all the data in the time course. *gpr-1/gpr-2* knockdown (KD) delayed spindle elongation. Control, n = 6; KD n = 8. (**B)** Snapshots of the curvature along the major axis of the cell. 0 denotes the cell center. Each curvature was extracted as in Fig. 5B. The control showed a clear peak at t = 400 s, while the KD showed a peak around t = 500 s. **(C)** Time courses of the solidity of the cell boundary. t = 0 s was defined as in (A). Consistent with (B), the control cells showed a clear drop in solidity before t = 400 s, whereas solidity remained high in KD until at least t = 500 s.

The cell curvature distribution along the major axis of the cell at different time points (t = 0 s corresponds to the nuclear envelope breakdown (NEBD) is shown in Figure 6B. Note that the *C. elegans* nuclear envelope disassembles very late (in mid-late anaphases) compared with those of vertebrates and *Drosophila* [28]. In the control cells, curvature distribution was initially almost uniform, with small negative values at both poles. Then, a prominent increase in curvature appeared at the center around t = 400 s, corresponding to the concavity in cytokinesis. *gpr-1*/*gpr-2* knockdown showed curvature snapshots similar to the control, although the appearance of a prominent concavity was delayed until t = 600 s. Note that the *gpr-1/gpr-2* knockdown is known to cause a positional shift of the dividing plane compared to the control [26]; our result is consistent with this report by showing the positional difference of the peak curvature at t = 500 and 600 s in gpr-1/gpr-1 compared to the control.

The time courses of the distance between spindle poles and solidity (see Methods for definition) showed that the cell morphology change followed the start of the increase in the distance between centrosomes (Fig. 6AC). Namely, a clear increase in the distance at t = 200 s (Fig. 6A) preceded the drop in solidity at t = 300 s (Fig. 6C) in the control. Quantitative image analysis showed consistency in cell morphology and spindle pole movement between empirical observation using time-lapse images and our simulation, subject to spindle elongation perturbation.

### Simulation of asymmetric cell shape

Next, we asked whether the asymmetric cell shape would affect the position of the cell division plane by modulating cytoplasmic flow. To compare the simulation results with the experimental data on cell morphology obtained by using a PDMS microchamber with sea urchin eggs [29], we set a triangular shape for the simulated cell (Fig. 7). The ratio of the base and the height of the triangle was set to 1:2, keeping the volume of the cell equal to that in the simulation showed in Figure 2–5. The triangular cell showed cortical cytoplasmic flow toward the cell center, similar to the ordinary round cell (Fig. 7B). Following cytoplasmic flow, a cell surface concavity appeared at the center of the spindle (Fig. 7C). This simulation reproduced the experimental data [29] that the cell division plane is located at the center of the spindle, although in our simulation the furrow position was very slightly shifted toward the narrow side of the cell (Fig. 7D). Thus, our computational model showed that the generation of the concavity is independent of cell shape.

**Fig 7.**
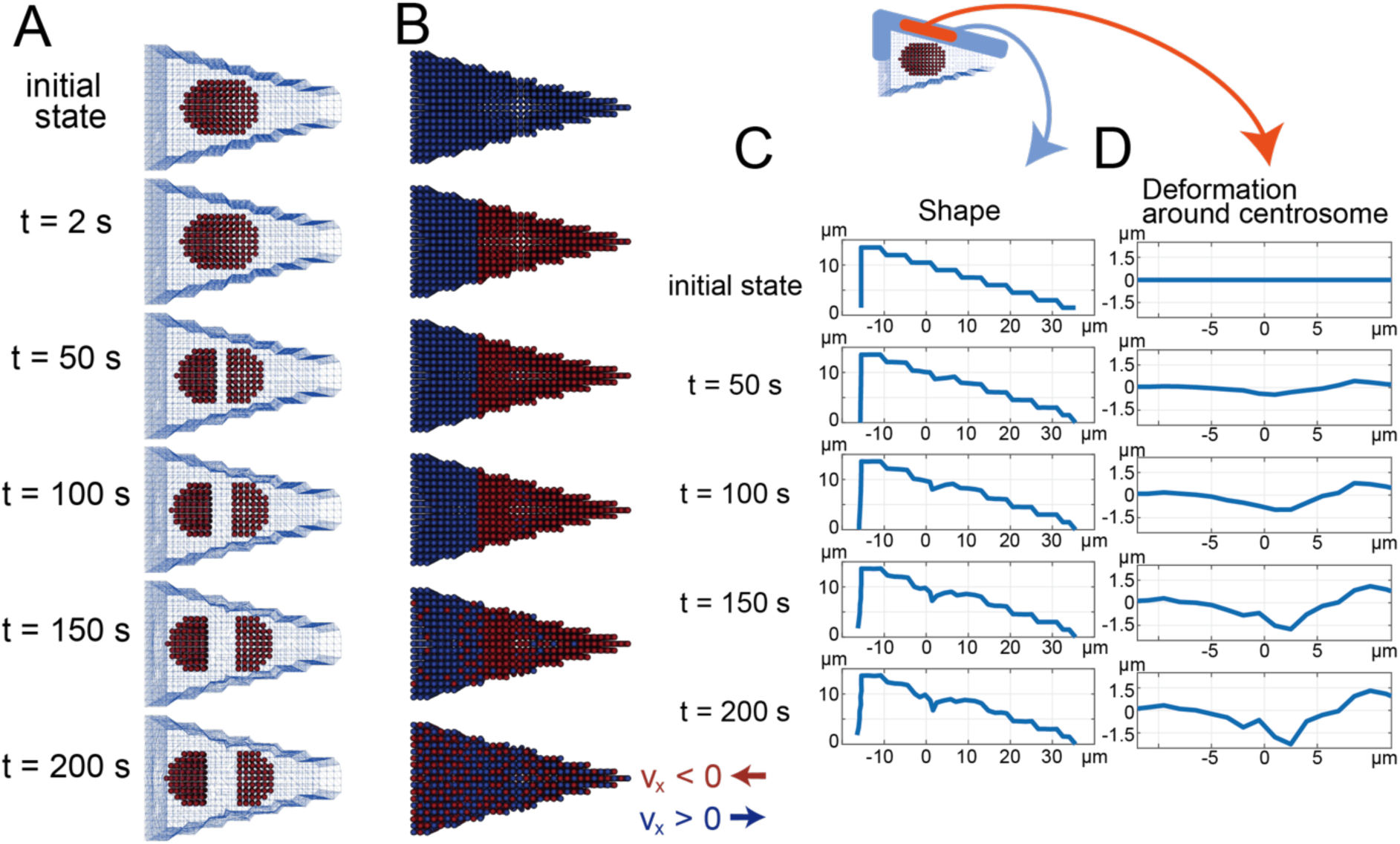
Simulation of a triangular cell. **(A)** Snapshots of the simulated cell shape. Red particles, centrosomes. **(B)** Simulated cytoplasmic velocity on the cell cortex. Color coding is as in Fig. 2. The triangular shape does not affect the distribution of velocity direction. **(C)** Cell morphology changes during simulation. The *xy* coordinates of the lateral cortex particles at the center of the cell along *z* = 0 are visualized to clarify morphological changes during the simulation. 0 denotes the initial *xy* position of the centrosome center. **(D)** Cortical deformation around the centrosome calculated by subtracting *y* coordinates from the initial position. 0 on the *x-*axis denotes the initial *x* position of the centrosome center. The *x*-position of the emerging concavity is almost 0, but is slightly shifted to the narrow side of the cell.

## Discussion

In this study, we constructed a mathematical model of the rheological interaction between cytoplasmic flow and cell cortex elasticity. Our model provides a simulation platform for exploring changes in cell morphology induced by cytoplasmic flow. Fluid dynamics associated with cytoplasmic activities follow the Navier–Stokes equations and the continuity equation [15]. The application of the strict MPS method to the simulation of cytoplasmic flow allowed us to quantitatively predict the dynamics of the cell’s internal pressure distribution, which induced a morphological change of the cell cortex, assuming the cell cortex is an elastic shell. This cortical deformation, in turn, affected fluid dynamics, creating a bidirectional mechanical coupling. Particularly, when the boundary condition changes dynamically, particle-based methods, including MPS, are a suitable simulation framework to predict the rheology of the material [30]. Our model provides insight into the interaction between the cell cortex and internal fluid dynamics. Time-lapse imaging of first embryonic cell division in *C. elegans* were consistent with our simulation that a reduction in the speed of spindle elongation delays the onset of an invagination on the cell surface. The consistency of quantitative image analysis and our simulation illustrates that despite the change in the parameters of the model components, the proposed model successfully reconstructed the outcome of physical interactions among spindle elongation, cytoplasmic flow, and the elastic cell cortex.

As a specific target, we focused on the determination of the position of the cell division plane in relation to cytoplasmic flow. Cytoplasmic flow has been observed during early cell cycle period and the effect on nuclear transport is proposed [31], but the fluid dynamic effect on cell shape remains unknown. Using our simulation model, we reproduced the generation of cytoplasmic flow by the elongation of spindle microtubules, as previously reported [15,32]. Our simulation quantitatively showed an immediate formation of the internal pressure distribution (Fig. 3). The change in cell surface pressure induced morphological changes of the cell cortex in our model. At the cell surface, flow from the two poles of the mitotic spindle collapsed in the middle of the spindle, and the invagination of the cell surface was induced at that position. In our simulation, the invagination was sustained, and the elasticity of the cell cortex caused cell concavity with a delay. The generated cleavage remained after the pressure distribution became uniform (Fig. 4). These observations have at least two implications for the mechanism that determines the position of the cell division plane. The collapse of the flow at the cell surface can help molecules required for cell division to accumulate at the cleavage furrow. In addition, the induction of the invagination at the furrow position may provide an initial mechanical cue for the formation of the cleavage furrow, before cell division.

The coupling between spindle elongation and the onset of furrow invagination was supported by our experimental data and the corresponding simulation. When we experimentally slowed down spindle elongation, the onset of the invagination was delayed relative to the onset of spindle elongation (Fig. 6C). In our simulation, slowing of spindle elongation delayed the onset of the cleavage furrow, without changing its position (Fig. 5AC at t = 150 s). Our simulation could also reproduce the cleavage furrow positions in cells of various shapes (Fig. 7), which is particularly advantageous in developmental studies. Overall, our simulation supports the hypothesis that the interplay between pressure distribution induced by cytoplasmic flow and cell cortex elasticity, but not the initial cell morphology, determines the position of the cell division plane.

Understanding the mechanistic nature of cytokinesis is complementary to molecular cell biology that has been actively pursued and is essential to understanding cell division. Previous models of furrow formation focused mainly on the molecular mechanisms of microtubules and chromosomes, and did not consider the effects of cytoplasmic flow and cell cortex pressure [33]. Each model refers to a specific organism and conditions, and predictions do not work under varying conditions. Our proposed model aims to describe the mechanical properties of cell division by focusing on the observable dynamics of the cell cortex, microtubules, and chromosomes.

An integrated view of our model and the previous models of cell division clarifies the perspective of how we understand this dynamic process composed of multiple highly interactive elements. The equatorial astral stimulation model proposes that astral microtubules deliver a positive signal to the cortex at the cell equator, promoting cytokinesis [4,32]. Our model extends this framework by incorporating cytoplasmic flow, which conveys positive cytokinetic signals to the equator and generates pressure that pushes the cell surface toward the astral microtubules. In the polar relaxation model, the cell equator maintains a constant cortical tension, whereas the daughter cell pole or body relaxes, allowing the cell to cleave [34]. Our simulation suggests that cytoplasmic flow in the polar region of the cell cortex promotes polar relaxation toward the cell equator. If we consider the cell cortex plane, cytoplasmic flow generates expansion in the polar region of the cell cortex whereas condensation occurs along the equator; from the viewpoint of the direction perpendicular to the cell cortex, outward pressure in the polar region and downward pressure at the equator correspond to cortical tension. This interpretation makes cytoplasmic flow as actual observable measurements for the polar relaxation model. The spindle midzone model proposed that continuous interaction between midzone microtubule bundles, which are structural signature of a central spindle, and the cortex is required for successful cleavage in tissue culture cells [35,36]. Our simulation supports the hypothesis that the central spindle plays a central role in the determination of the dividing position, and the implicit mechanism postulates the localization of positive factors for cytokinesis is caused by spindle microtubules, similar to the equatorial stimulation model. The transport role of the microtubules is independent of their mechanical properties proposed by our study; therefore, both mechanisms may serve cooperatively or one of them may prevail.

## Materials and Methods

### Strains and RNAi

Wild-type N2 Bristol worms were obtained from the *Caenorhabditis* Genetics Center (CGC). All worms were hermaphrodites and were cultivated on OP50 bacterial food using standard techniques [37]. CAL0092 *unc-119*(*ed3*) III; *ddIs6*[*tbg-1*::GFP + *unc-119*(+)]; *weIs15*[*unc-119*(+); *pie-1p*::GFP::FYVEx2] was described in [38].

The template for RNAi of *gpr-1*/*gpr-2* was amplified by PCR from genomic DNA with primers based on sequences in the Phenobank database (http://worm.mpi-cbg.de/phenobank2) [39]. double-stranded RNA was derived from in vitro transcription using the template. RNAi was performed by injecting the dsRNA into adult *C. elegans* worms, as previously described [40].

### Spindle elongation model

We postulated that spindle elongation is driven by motor proteins (such as dynein and kinesin on microtubules) independently of the cytoplasmic flow or change in cell morphology, and we gave typical elongation patterns to the simulator. Such a pattern is shown in Figure 2; it was based on the measurements of the distance between centrosomes in fluorescence time-lapse images. The simulation parameters are listed in Table 1; they are biologically appropriate values from published papers. The time course of spindle elongation was given as an input and was used by the model to compute other physical values.

### Cell morphology model

On the basis of the simulation model for the morphology of red blood cells [41], we defined the cell surface particle as a triangular elastic mesh by the following equations. The elongation spring force F_s_ and the bending spring force F_b_ were given as follows:

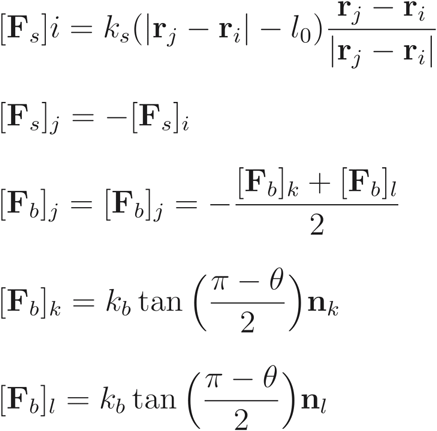

where *i*, *j*, *k*, and *l* denote the indices of the four particles (Fig. 8), r_i_ denotes the effective radius for particle *i*, and r*_j_* - r*_i_* denotes the distance between particles *i* and *j*. *l*_0_ denotes the initial average distance between cell surface particles, k_s_ is a spring constant for the extension axis, θ is the angle defined with the neighboring particles, *k_b_* is the spring constant for the bending axis, and n*_k_* and n*_l_* are the perpendicular vectors for the triangles ikj and ijl, respectively.

**Fig 8.**
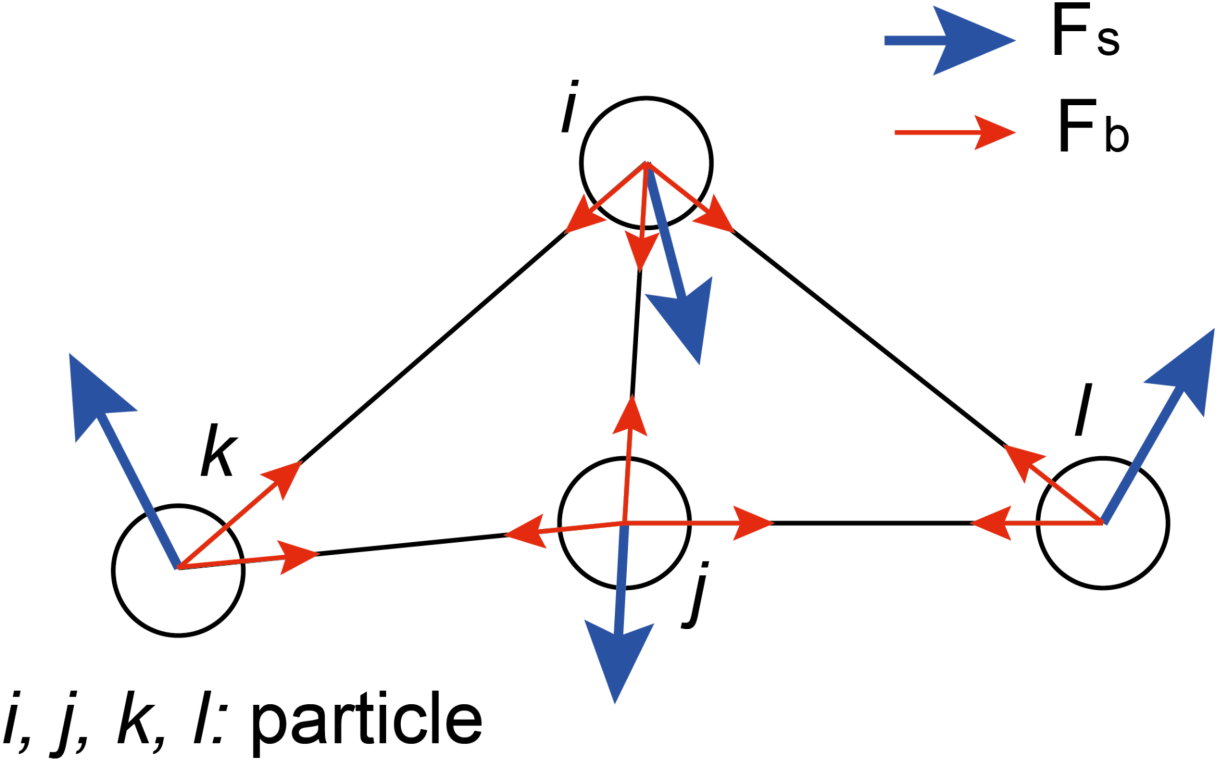
Cell surface mesh particle model. Each particle is connected to the neighboring particle to form a triangular mesh. Elongation spring force (F_s_) and bending spring force (F_b_) were calculated on the basis of the defined mesh during simulation and reflect cell surface morphology. The redrawn spring model is based on [21].

To generate the triangular mesh, we used the Point Cloud Library (PCL) [42], which consists of a set of cell surface particles. PCL is specialized on 2D/3D images and point cloud processing and includes many algorithms such as filtering, feature estimation, surface reconstruction, and segmentation. We used the Greedy Projection Triangulation function to reconstruct the surface from point clouds. The procedure is as follows.

1. Export the coordinates (*x*, *y*, *z*) of the cell surface particles to a file from the simulator.
2. Transform the exported coordinates into the Point Cloud Data (PCD) format.
3. Input the list of coordinates in PCD format to Greedy Projection Triangulation and output in Visualization Toolkit (VTK) format.
4. Input the VTK file into the simulator and load the relationship between each particle based on the original coordinates.

The simulator computes the force on the cell surface on the basis of the relationship between the particles and conveys it to the external force term of the Navier–Stokes equation.

### Cytoplasmic flow simulation using the MPS method

We implemented the simulation of the cytoplasmic flow according to the previous work for a *C*. *elegans* embryo [15]. Cytoplasmic flow is defined as fluid following the Navier–Stokes equation:

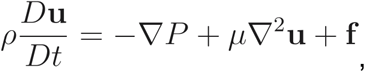

where *D*u/*Dt* denotes the Lagrangian derivative, u denotes the velocity vector for a particle, ρ denotes the density, P denotes the pressure, μ denotes the viscosity, and f denotes the external force.

To compute the unknown pressure P, we considered the equation of continuity, which represents the conservation law for fluids and the incompressible condition for the intercellular environment. Then, we derived the following equations:

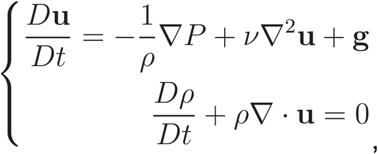

where ν represents the kinematic viscosity coefficient derived from μ/ρ. We described the position of the particle *i* at the step *k* as r_i_*^k^*, and then the next position r_i_^k+1^ is calculated from the governing equation and velocity u*^k^*^+1^_i_ as

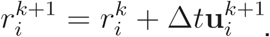

#### MPS method

We implemented the above governing equations of continuity and the Navier–Stokes with the particle method [30], in which fluid is discretized into arbitrarily sized lumps, which are then considered as particles having variables corresponding to physical properties. We selected the MPS algorithm for incompressible fluids, in which particle density n*_i_* of particle *i* is calculated as:

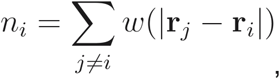

where r*_i_* denotes the positional vector of particle *i*, and n_i_ is the sum of weighted distances among particles in an effective distance. The weight *w* is usually defined as

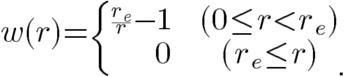

However, because this definition makes the pressure anomaly high in our case, we used the following equation for *w* [41] to make the computation stable:

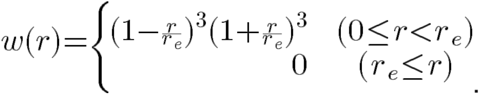

In particle simulations, the initial uniform particle density n° is defined as a calibration value to compute the degree of fluid condensation. The change in fluid density is approximated as follows:

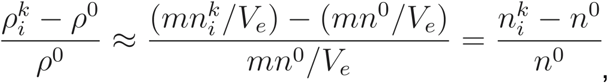

where ρ^0^ denotes the standard density of the fluid and ρ^k^_i_ denotes the fluid density for particle *i* at the time step *k* under the given pressure. The constant *m* denotes the mass of one particle, and V_e_ is the volume inside the effective radius. From this equation, the degree of fluid condensation can be evaluated only from the particle density. The MPS method uses the following steps.

Step 1. Load initial values and parameters

Step 2. Compute the tentative velocity and position on the basis of the viscosity term, then move the particles

Step 3. Compute the tentative particle density

Step 4. Compute the pressure using the pressure Poisson equation

Step 5. Compute the velocity correction from the pressure gradient and apply it to update the particle velocity

Step 6. Exit decision (or back to step 2).

#### Implicit method

Fluid simulation for micrometer order often becomes unstable, and numerical values diverge with explicit calculation. Therefore, we used a fully implicit method in which both the pressure gradient and viscosity terms are solved implicitly, unlike the conventional semi-implicit method where only the pressure gradient is treated implicitly. Although our approach increases the computation time, the computational outcome becomes stable because updating time dependent values are free from a constraint for Δt. The particle *i* of the tentative velocity at time step *k* for the explicit and implicit methods are as follows, respectively:

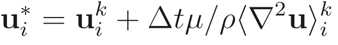

and

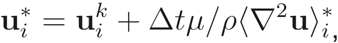

describing a viscosity as the second terms of the right-hand side of the equations.

The explicit method uses only the known values, while the implicit method uses an unknown value in the viscosity term.

The following Laplacian model is used to discretize the Laplacian [30]:

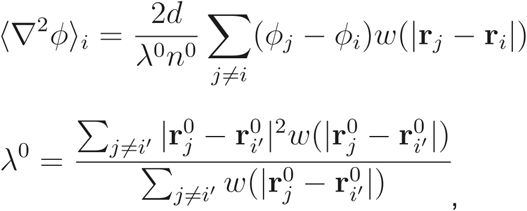

where λ^0^ denotes the weighted average of the squared distance between particles within an effective radius. 2*d*/λ^0^*n*^0^ is the constant for adjusting the diffusion speed; in other words, the fitting constant for adjusting the increase of the variance in one time step to the analytical value. The expanded form of u*_i_ based on the above Laplacian model, becomes as follows:

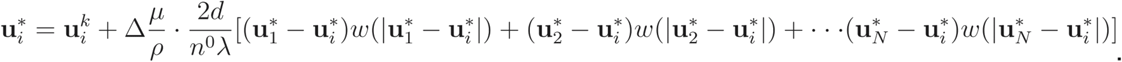

Replacing the terms in the above equation for simplification as

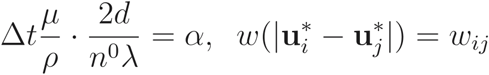

generates the following simultaneous equations for each particle *i*:

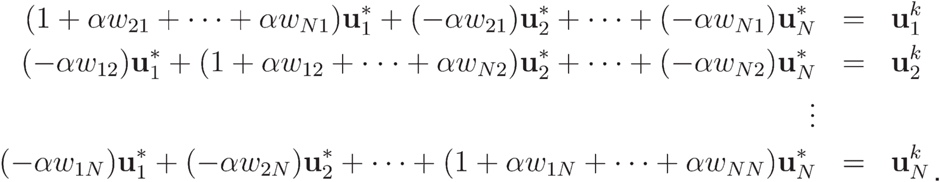

By describing the set of simultaneous equations for *a_ij_* and *b_i_* as

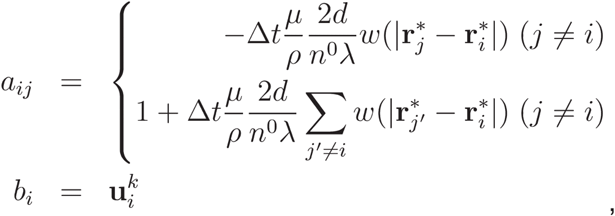

we obtain a matrix representation of the simultaneous linear equations. Then, we solved the equation numerically using Gauss elimination and conjugate gradient methods:

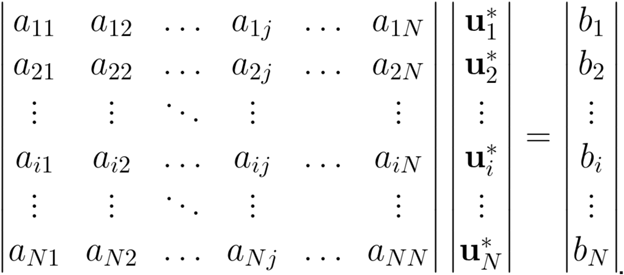

### Boundary conditions

The boundary condition was given below to solve the Poisson equation for pressure. The pressure of the particles on the free surface (liquid surface) was fixed to 0 Pa as a Dirichlet boundary condition. If particle *i* satisfies the n_i_ < βn^0^ condition, it is defined to be on a free surface. We used β = 0.97 as it is generally used for fluid simulation.

The Neumann boundary condition was used for the wall. Since the wall does not transmit fluid particles, two layers of wall particles holding a pressure value were aligned at the border of the fluid and the wall; additionally, multilayers of dummy particles, which do not have a pressure value, were aligned outside the wall particles. These dummy particles approximately express the Neumann boundary condition with zero pressure gradient.

For velocity, the no-slip condition was given at the wall surface by setting it at 0 m/s in viscosity calculations. If no strict no-slip condition is necessary, the velocity of the wall particle was fixed at 0 m/s.

### Strategies for reducing computation time

We used three strategies: the conjugate gradient method, parallel computing using OpenMP, and the bucket data structure using a linked list. Our particle simulator with an implicit method needs ≤4 computations to solve the equations with viscosity (*x*, *y*, *z* directions) and the pressure gradient term in one time step. The runtime for solving linear simultaneous equations is proportional to the size of the coefficient matrix for these equations, which is N × N (N, number of particles in the simulated space). Because solving simultaneous equations is the bottleneck of the simulation runtime with the particle method, the selection of an efficient algorithm for solving these equations drastically reduces the runtime. In this study, N ≈ 20,000. Among many algorithms, we used the conjugate gradient algorithm [43] because it leverages the characteristics of the coefficient matrix in the particle method through an iterative procedure. At the implementation level, we used OpenMP to take advantage of parallel computation [44]. In the codes, for loops are computed in parallel by calling the OpenMP directives through including a library file omp.h in compiling. We used the bucket data structure to effectively explore neighboring particles. MPS requires calculations for the interaction among particles within the effective radius. When judging whether the distance between the target particles is smaller than the influential radius, N × N calculations based on all particle combinations are required. To reduce the computation time, we followed the method described in Harada et al. [45] for filtering closely neighboring particles using the bucket method.

All code written in support of this publication is publicly available at https://github.com/funalab/CeMPS

### Quantitative image analysis

Embryos were imaged as described [46] with the following modifications. Embryos were removed from adult worms and mounted on a microscope slide. Fluorescence signals excited with a 488 nm laser were visualized using a spinning-disk confocal system (CSU-X1; Yokogawa, Tokyo, Japan) mounted on an inverted microscope (IX71; Olympus, Tokyo, Japan) equipped with a 100×, 1.40 N.A. objective lens (UPLSAPO; Olympus). Digital images of control and *gpr-1*/*gpr-2* (RNAi) embryos were acquired every 2 s with an EM-CCD camera (iXon; Andor Technology, Belfast, UK) controlled by Metamorph software (Molecular Devices, CA, USA) with an exposure time of 50 ms.

We measured the positions of centrosomes using the manual tracking tool in the open-source image analysis software Fiji [47]. The time courses of the distance between the centrosomes were subtracted from the initial distance of each time course to estimate the elongation of the distance between centrosomes in different cells. The initial time point (t = 0 s) was defined by the frame that captured the NEBD. To quantify cell morphology, we defined the curvature *κ* of the cell edge extracted from a fluorescence image as:

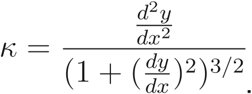

Following a previous study that calculated the cell curvature [48], we used Fiji to extract the cell region. To calculate the curvature, the first- and second-order derivatives were obtained numerically using the pixel difference.

We also used solidity for the morphological features of the entire cell. Solidity is defined as the area of a cell divided by its convex hull area, and reflects the extent of the total concavity of the cell.

## Acknowledgements

We thank Kazuko Oishi for technical assistance with imaging of *C. elegans* embryos. This work was supported by JSPS KAKENHI Grant Number JP15KT0083.

